# Persistent fibrosis and decreased cardiac function following cardiac injury in the *Ctenopharyngodon idella* (grass carp)

**DOI:** 10.1101/627752

**Authors:** Daniel W Long, Charles H Webb, Yadong Wang

## Abstract

Following the discovery of heart regeneration in zebrafish, several more species within the Cyprinidae family have been found to have the same capability, suggesting heart regeneration may be conserved within this family. Although gonad regeneration has been observed in grass carp (*Ctenopharyngodon idella*), one of the largest cyprinid fish, the species’ response to cardiac injury has not been characterized. Surprisingly, we found cardiomyocytes do not repopulate the injured region following cryoinjury to the ventricle, instead exhibiting unresolved fibrosis and decreased cardiac function that persists for the 8-week duration of this study. Compared to other cyprinid fish studied, infiltration of macrophages is delayed and muted in this model. Additionally, fibroblasts are depleted following injury, a phenomenon not previously described in any cardiac model. This study shows that heart regeneration is not conserved among the Cyprinidae family and suggests the important role of non-fibroblasts in chronic fibrosis. Further study of these phenomenon may reveal the underlying differences between regeneration versus unresolved fibrosis in heart disease.

**Summary statement:** Grass carp, a member of the Cyprinidae family that includes regenerative zebrafish, do not regenerate functional cardiac tissue after cryoinjury. Instead, healing progresses through collagen deposition and scar formation.

## 1. Introduction

Organ regeneration is a phenomenon that occurs in select species across the animal kingdom (Lenhoff and Lenhoff, 1986; Morgan, 1901). Humans are generally incapable of regenerating damaged tissues with a few notable exceptions (Michalopoulos, 2007). Heart regeneration specifically is not seen in humans but is reported in several adult non-mammalian species and neonatal mammals (Cano-Martínez et al., 2010; Liao et al., 2017; Porrello et al., 2011; Poss et al., 2002; Ye et al., 2018). Following the discovery of cardiac regeneration in zebrafish after ventricle resection in 2002 (Poss et al., 2002), several varieties of fish have been investigated for their regenerative potential after cardiac injury. To date, cardiac regeneration has been observed in several cyprinid fish, including the zebrafish, giant danio, and goldfish, along with a surface variety of *Astyanax mexicanus* outside the Cyprinidae family (Grivas et al., 2014; Lafontant et al., 2012a; Poss et al., 2002; Stockdale et al., 2018). In contrast, the cave-dwelling Pachón variety of *Astyanax mexicanus* and medaka (*Oryzias latipes*) do not exhibit signs of cardiac regeneration following injury, neither of which are in the Cyprinidae family (Fig. 1A) (Ito et al., 2014; Stockdale et al., 2018).

**Figure 1:**
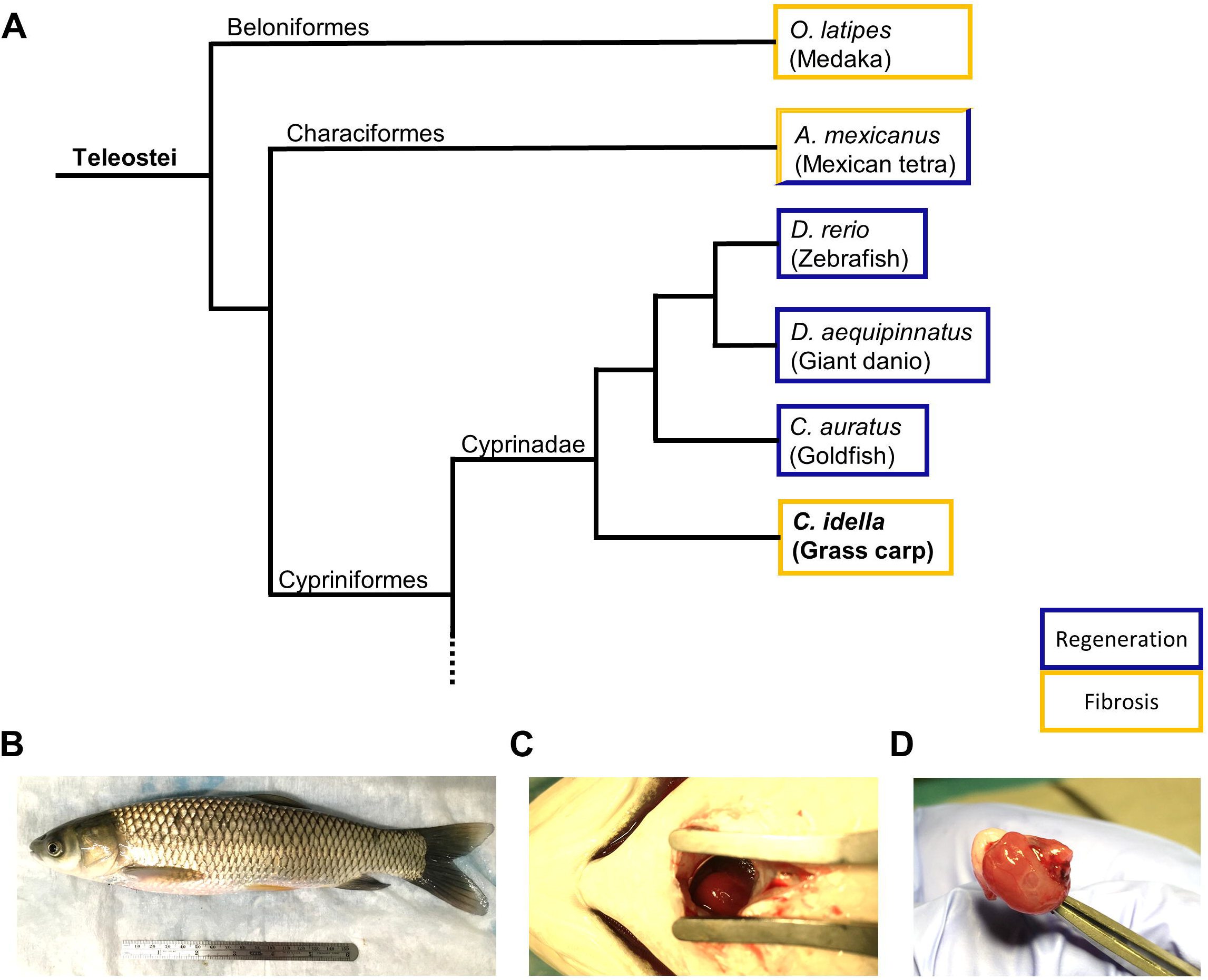
Grass carp characterization and taxonomy. (A) All previously investigated cyprinids exhibit cardiac regeneration following injury with exception to grass carp presented in the present study. Medaka and at least one variant of *A. mexicanus*, distant relatives of the grass carp, instead heal via fibrosis following cardiac injury. (B) Grass carp used in this study were approximately 300mm in length (total length) and 435 ± 112 g in mass. (C) Alm retractors were used to separate the operculum and allow direct visualization of the ventricle. The injured zone at the apex is outlines in yellow. (D) Explant at 56 DPI reveals white scar tissue rather than healthy pink tissue indicative of full regeneration.

While cardiomyocyte proliferation was thought to be the main driver of cardiac regeneration in fish, the recent discovery of widespread cardiomyocyte proliferation in non-regenerative Pachón *Axtyanax mexicanus* indicates that there are alternative regulators of cardiac injury at play (Stockdale et al., 2018). However, with only two non-regenerative fish species currently reported, it is difficult to assess which mechanisms are essential regulators of regeneration versus chronic fibrosis. Therefore, research on responses to cardiac injury in additional fish species will inform what cellular behaviors are chiefly responsible for driving cardiac regeneration versus chronic fibrosis. Grass carp, or *Ctenopharyngodon idella*, are an Asian variety of carp within the Cyprinidae family of fish (Fig. 1A). Unlike previously reported zebrafish (<1 g) (Gómez-Requeni et al., 2010), giant danio (1.6 g) (Lafontant et al., 2012b), medaka (<2 g) (Ding et al., 2010; Lafontant et al., 2012a), and goldfish (3.4 g) (Grivas et al., 2014) models of cardiac injury, grass carp used in this study are over 400 g and over 30 cm in length (Fig. 1B). These fish are capable of exceeding 25 kg in mass and 1 meter in length in aquaculture conditions, thus their size is several orders-of-magnitude greater than the previously studied species (Stich et al., 2013). As an invasive species to the United States, early efforts to sterilize these fish by removal of the gonads were ineffective due to their ability to regenerate reproductive organs within months of removal (Underwood et al., 1986). The regenerative potential of other organs in this fish have not been further explored. In fact, the regenerative potential of cardiac tissue in any large fish species is unknown.

In the present study we design a method to reproducibly injure the heart of the grass carp, with a survival rate >95% 24 hours post-surgery. We investigated the cellular and functional response of the heart following injury. Our study provides a surgical model for investigation of cardiac injury in large fish species and suggests that grass carp are the first reported cyprinids that exhibit chronic fibrosis rather than functional regeneration. Additionally, fibroblast and macrophage responses to injury are different from reported cyprinid fish species and warrant further investigation.

## 2. Results

### 2.1 Cardiac wound healing following cryoinjury in the grass carp

The use of cryoinjury was selected over resection due to limited access to the apex in grass carp heart with scissors (Fig. 1C). The use of a 2 mm stainless steel cryoprobe resulted in an injury covering 10-15% of the ventricle area at its maximum depth (Fig. 2B), similar to previously reported goldfish injury (Grivas et al., 2014). Although this is a relatively small injury compared to standard zebrafish and giant danio models (Chablais et al., 2011; Lafontant et al., 2012a), larger injuries resulting from longer freezing times and larger probes resulted in greater than 85% mortality within 14 days of injury. Acid fuchsin orange G (AFOG) staining of the fish ventricle was used to identify and quantify the injury size over 8 weeks post-injury. Uninjured hearts contained a collagen-rich bulbus arteriosus in blue and a healthy ventricle and atrium stained orange (Fig. 2A). Fibrin deposits and the injury site were easily visible by 1 day post injury (DPI), which was converted to a collagen-rich scar by 56 DPI (Fig. 2B,C). Quantification of the injury size over time revealed that although the injury does reduce in size by 14 DPI relative to 1 and 4 DPI (p < 0.05 and p < 0.01, respectively), there is no further reduction between 14 and 56 DPI (Fig. 2D). Additionally, a white scar region is grossly visible on the ventricle 56 DPI at explant, indicating an incomplete regeneration of the injured ventricle (Fig. 1D).

**Figure 2:**
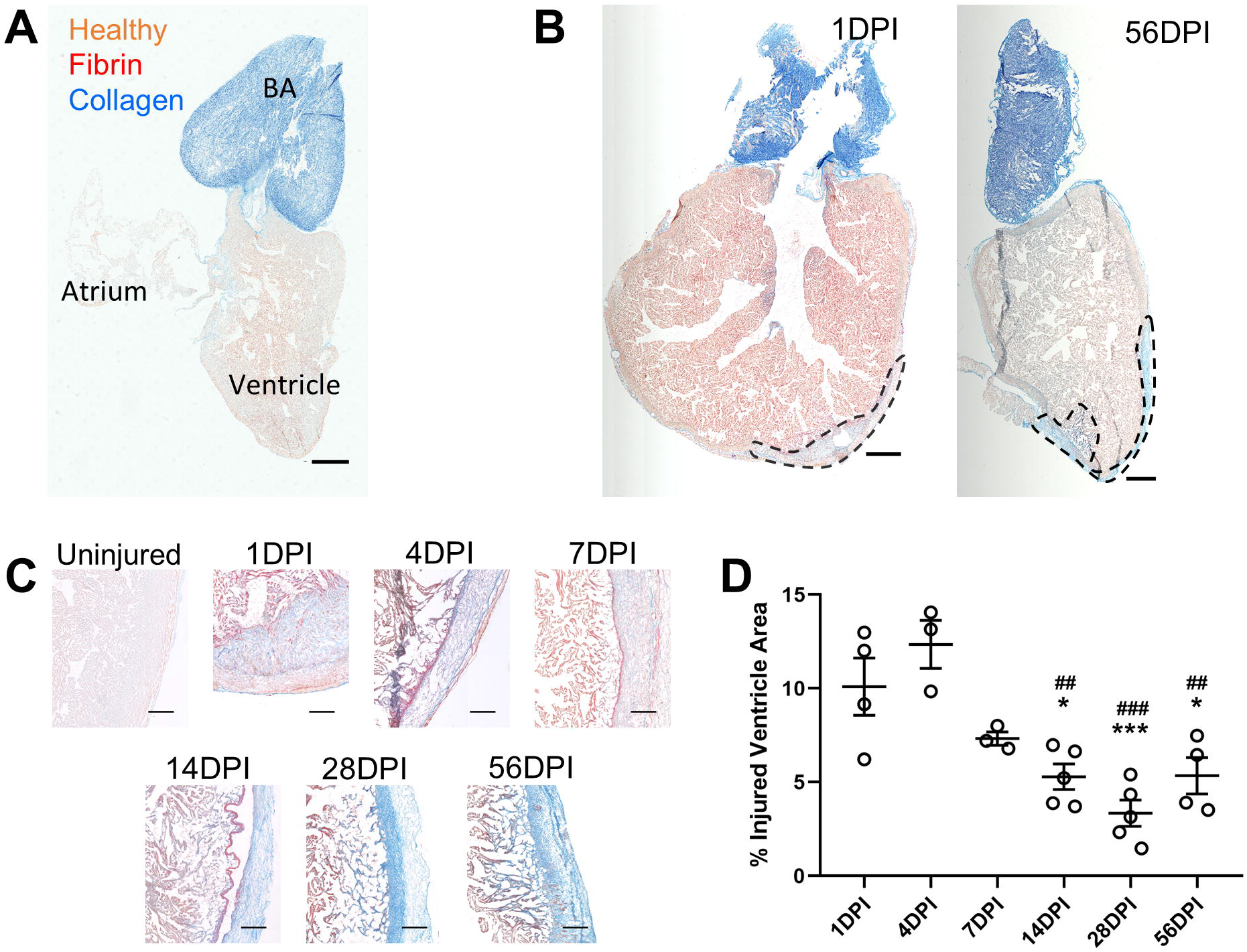
Heart anatomy and injury response. (A) AFOG-stained grass carp hearts reveal a single atrium, a ventricle composed of a cortical outer myocardium and trabecular inner myocardium, and the collagenous bulbus arteriosus (BA), which serves at the outflow tract (scale bar = 1 mm). (B) The injury site is visible at the ventricle apex 1 DPI, which is still visible 56 DPI as a fibrous scar, outlined in black (scale bar = 1 mm). (C) AFOG staining reveals the progression of the injury site from a fibrin-rich wound 1-7 DPI to a collagen-rich scar by 28-56 DPI (scale bar = 200 μm). (D) Quantification of the injury size reveals a significant reduction from early to late-stage injuries, although injury size does not decrease further after 14 DPI. (*p < 0.05, ***p < 0.001 relative to 1 DPI; ^##^p < 0.01, ^###^p < 0.001 relative to 4 DPI).

### 2.2 Cardiac function fails to recover following injury

In addition to histological assessments, echocardiography-based measurements were used to assess functional regeneration of cardiac tissue. B-mode echocardiography along the long-axis was used to measure ventricle area and length at systole and diastole (Fig. 3A). Cryoinjury to the hearts resulted in an immediate decrease in fractional area change (FAC) that persisted through the rest of the 56-day study period (Fig. 3B; p < 0.001 relative to uninjured at 4 DPI, p < 0.05 relative to uninjured at 1, 7, 14, and 56 DPI). Similarly, fractional shortening (FS) values were significantly reduced by 1 DPI and persisted through 56 DPI (Fig. 3C; p < 0.001 relative to uninjured at 1, 4, 7, 14, and 56 DPI). Interestingly, neither FAC nor FS values are significantly lower than uninjured fish at 28 DPI, although the scar is clearly still present (Fig. 2). This assessment paired with measurements of injury size (Fig. 2) indicates a lack of cardiac regeneration both histologically and functionally.

**Figure 3:**
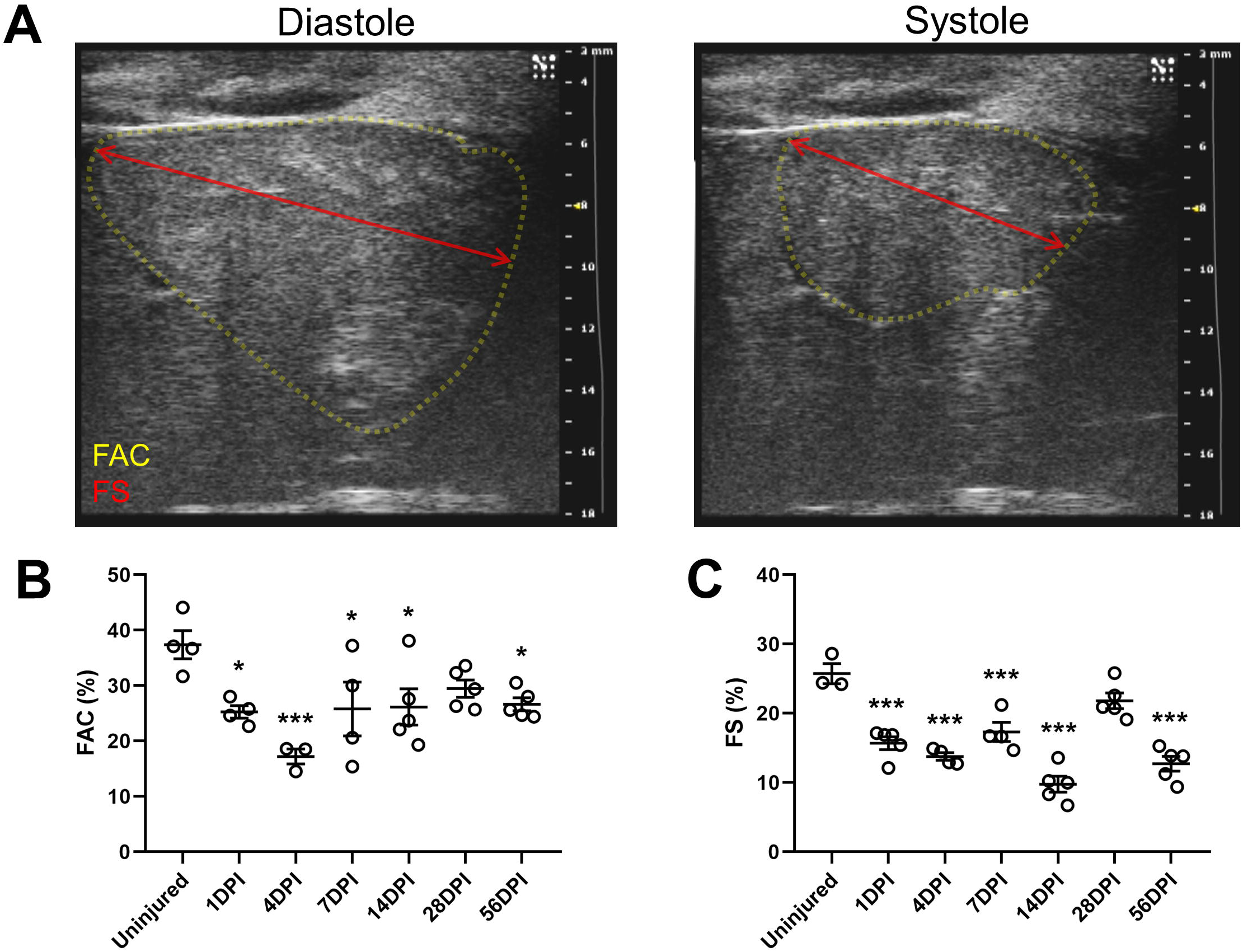
Cardiac function does not recover following injury. (A) Long-axis B-mode echocardiography of the heart at diastole and systole were used to quantify FAC and FS measurements following injury. (B) FAC values, though they improved from their minimum at 4 DPI, did not recover healthy function by 56 DPI. (C) Similarly, FS values did not improve to baseline values by 56 DPI, although FS at 28 DPI appeared to partially recover. (*p < 0.05, *** p<0.001 relative to uninjured).

### 2.3 Cryoinjury induces localized and acute apoptosis

Cardiac injury, especially cryoinjury, results in high levels of apoptosis in the injury zone in other models (Chablais et al., 2011; Darehzereshki et al., 2015; Polizzotti et al., 2015; Talman and Ruskoaho, 2016). TUNEL staining of cardiac tissues in the injured zone revealed high levels of apoptosis at 1 and 4 DPI (Fig. 4A,B; p < 0.001 relative to uninjured), which were back at baseline levels by 7 DPI. Except for a slight increase in TUNEL+ cells at 14 DPI (p < 0.05 relative to uninjured), apoptosis levels were low for the remainder of the study period. TUNEL staining in remote regions of the heart did not reveal a significant increase in apoptosis at any point after injury (Fig. 4A,C). Together these results indicate that cryoinjury in the grass carp heart causes an acute response of localized apoptosis that quickly resolves without spreading into otherwise healthy tissue.

**Figure 4.**
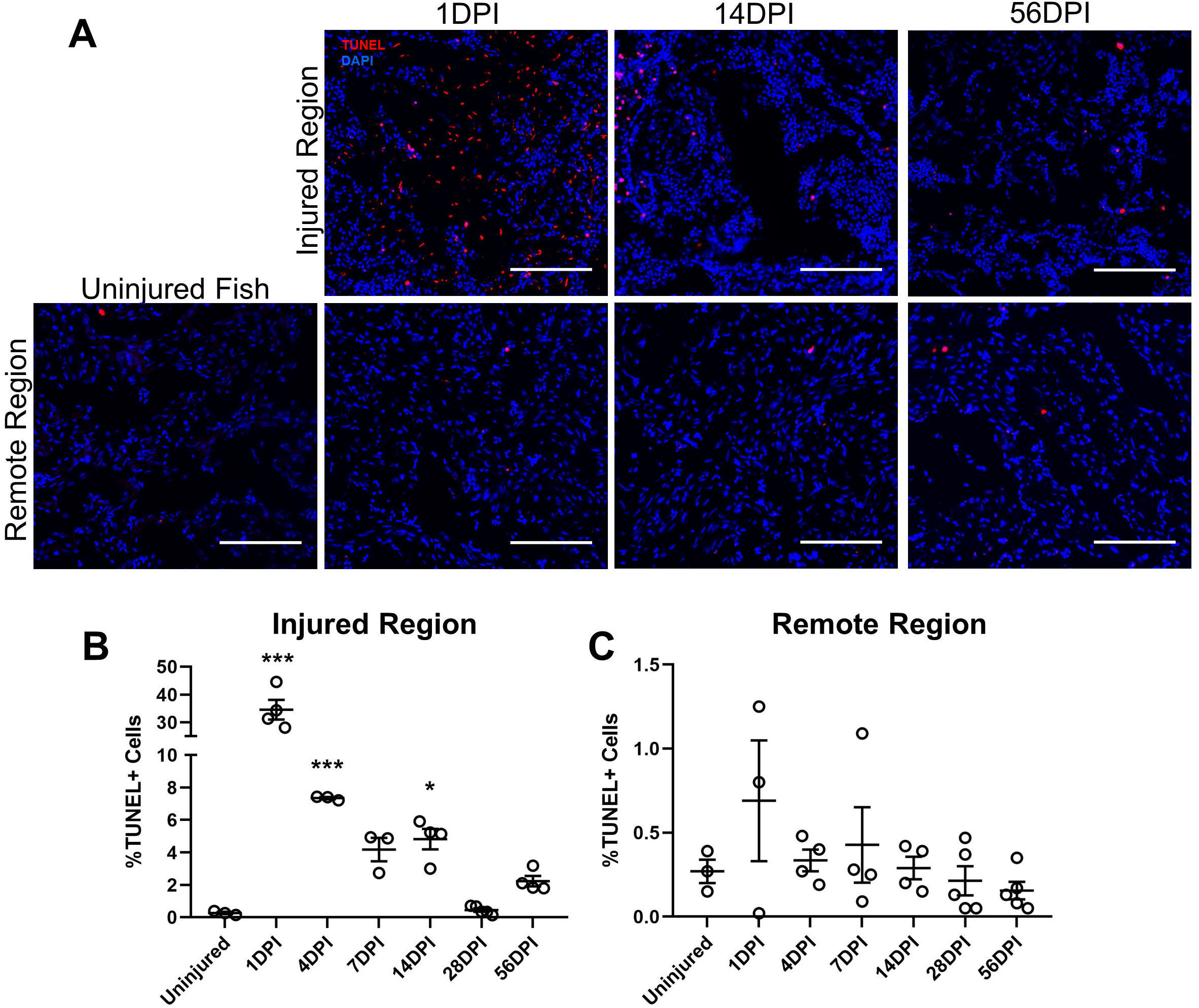
TUNEL Staining reveals a localized burst of apoptosis in the injured region. (A) Apoptotic cells are visible at all points after injury, although this is most pronounced in the injured region and at early timepoints following injury. (B) Quantification of TUNEL+ cells reveals high levels of apoptosis at 1, 4, and 14 DPI in the injured region. (C) TUNEL+ cells are not present in high quantities in remote regions of the injured heart at any point after injury. (scale bar = 100 μm; *p < 0.05, ***p < 0.001 relative to uninjured).

### 2.4 Cardiomyocyte proliferation is limited following cryoinjury

Proliferation of existing cardiomyocytes is a known driver of cardiac regeneration in zebrafish and neonatal mice (Chablais et al., 2011; Jopling et al., 2010; Porrello et al.; Schnabel et al., 2011). Staining for cardiomyocytes (cTnT) and phospho-histone H3 (pH3) revealed low to absent levels of cardiomyocyte proliferation at every timepoint following injury (Fig. 5A,B), which at least partially accounts for a failure of cardiomyocytes to repopulate the injured area (Fig. 5A). However, proliferation of non-myocytes was also quantified and revealed an increase in proliferative cells at 14 and 28 DPI specifically in the injured region (Fig. 5A,C; p < 0.01 at 14 DPI and p < 0.001 at 28 DPI relative to uninjured fish; p < 0.01 at 14 DPI and p < 0.05 at 28 DPI relative to remote regions at that timepoint). This indicates that the injured region is being repopulated by non-myocytes following cryoinjury. This could potentially also be due to proliferation of dedifferentiated cells that fail to fully differentiate back into functioning cardiomyocytes.

**Figure 5.**
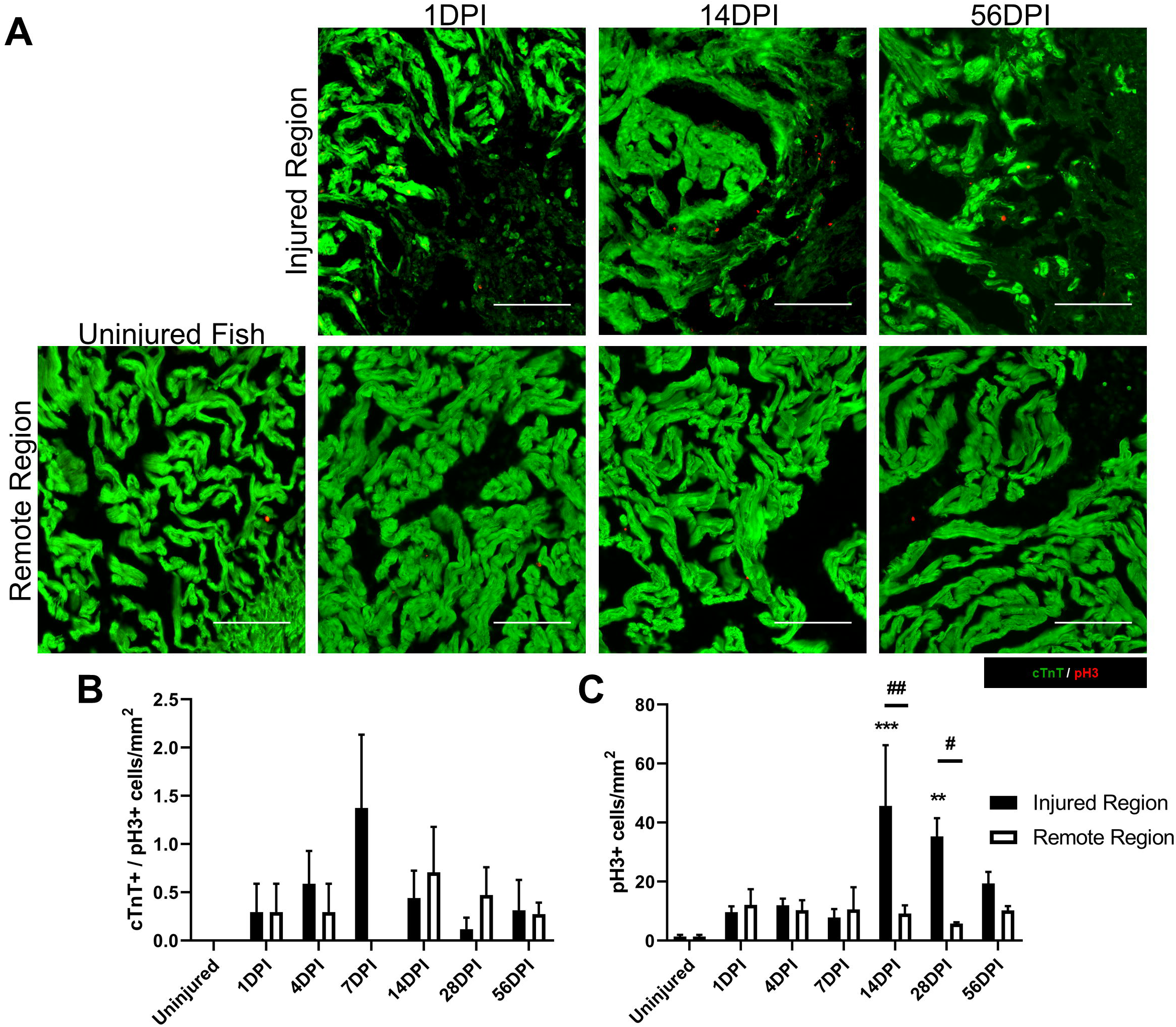
Local proliferation of non-cardiomyocytes contributes to fibrosis. (A) Co-staining of cTnT+ cardiomyocytes and pH3+ cells reveals an increase in non-myocyte proliferation at 14 DPI specifically in the injured zone. Quantification of proliferating cell types reveals that (B) cardiomyocytes do not proliferate to repopulate the injured region at any point after injury, but (C) a proliferating non-myocyte population within the injured zone is present at 14 and 28 DPI (scale bar = 100 μm; **p < 0.01, ***p < 0.001 relative to uninjured; #p < 0.05, ##p < 0.01 between regions).

### 2.5 Grass carp exhibit persistent and chronic fibrosis following injury

While fibrosis and cardiac regeneration are not mutually exclusive, scar resolution is necessary to allow infiltration of healthy tissue into the injured site. Masson’s trichrome staining revealed that scar formation, visible as collagen-rich blue regions, started by 14 DPI and persisted through 56 DPI (Fig. 6A). These samples were stained with picrosirius red to assess the maturation and alignment of collagen fibers within the scar (Fig. 6B). Scars at 14 DPI contained a large fraction of green fibers, indicative of immature collagens while scars at 28 and 56 DPI contained predominantly mature red fibers (Fig. 6B,C; mature fiber fraction p < 0.001 between 14 DPI and both 28 and 56 DPI). Collagen fibers in the scar were aligned along the ventricle wall at 14 DPI but became more randomly oriented by 56 DPI (Fig. 6B,D; p < 0.01 between 14 and 56 DPI. This correlates with an increased presence of collagen fibers extending form the ventricle surface into the trabecular myocardium. Together these data indicate that the scar tissue is maturing over time with no evidence of regression within 8 weeks.

**Figure 6:**
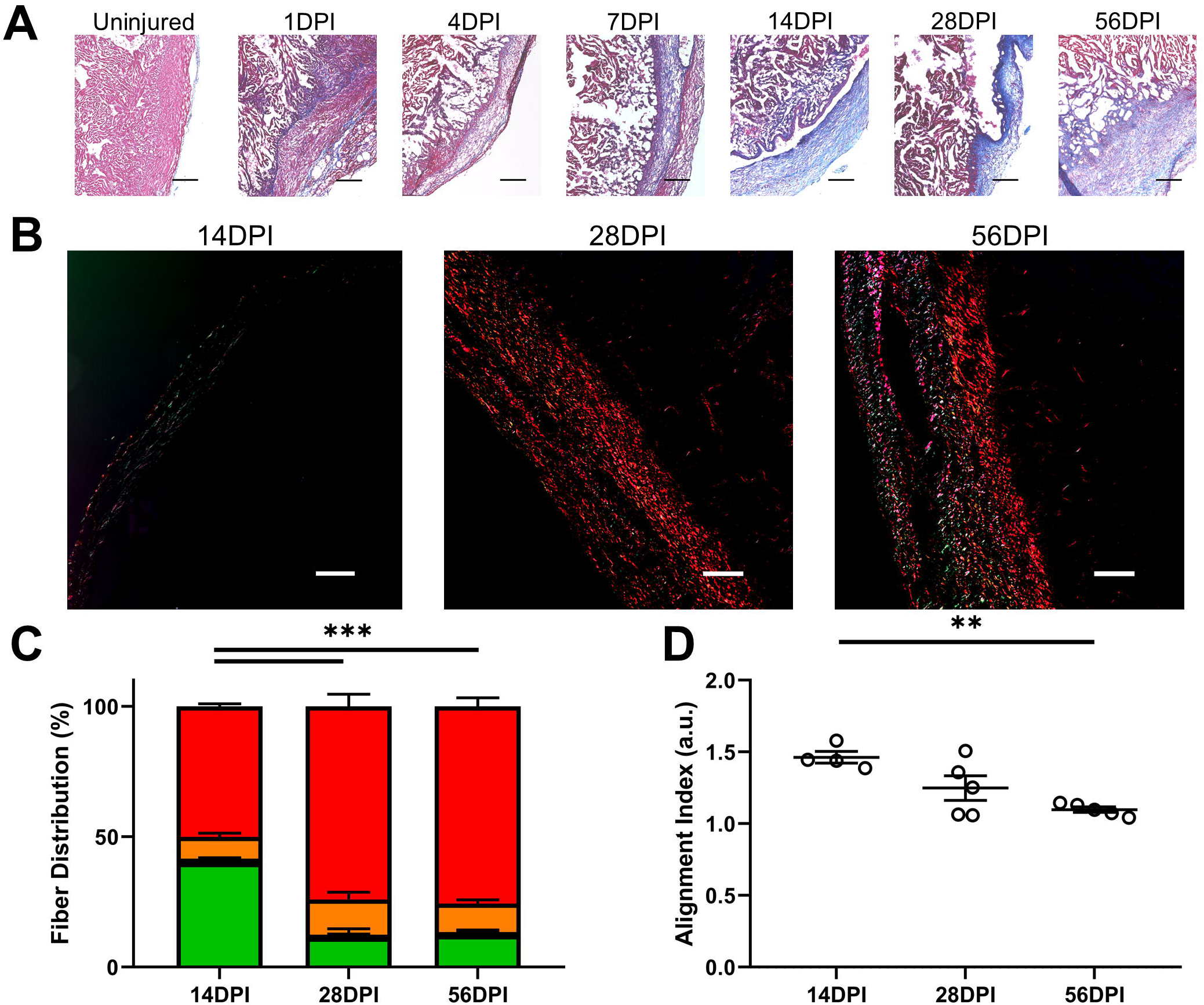
Scar formation and maturation in the ventricle following cryoinjury. (A) Masson’s trichrome staining reveals the formation of a fibrotic scar in blue visible at 14, 28, and 56 DPI (scale bars = 200 μm). (B) Picrosirius red staining of sections at the timepoints following scar formation show the presence of collagen fibers migrating into the intact myocardium (scale bars = 100 μm). (C) Analysis of picrosirius red images by fiber color shows a shift from immature green fibers at 14 DPI to predominantly red fibers at 28 and 56 DPI. (D) Collagen fibers become less aligned over time, resembling a cross-hatch pattern in the injured region. (**p < 0.01, ***p < 0.005 between groups).

### 2.6 Fibroblasts depletion occurs across the ventricle following injury

Conventional wisdom on fibrosis in mammals and fish studied so far suggests fibroblasts and activated myofibroblasts play a major role in ventricular remodeling, depositing and restructuring the extracellular matrix (ECM) within the injured region. Therefore, we quantified the density of vimentin+ fibroblasts in both injured and remote regions of grass carp hearts to assess their contribution to fibrosis. Unexpectedly, fibroblast density decreased in both injured and remote areas of the heart following cryoinjury (Fig. 7A). This depletion of vimentin+ fibroblasts extended throughout the 8-week study period in both injured and remote regions of the heart. Fibroblast density was lowest at 4 DPI in the injured region and 7 DPI in remote regions of the heart (Fig. 7C; injured region: p < 0.05 at 1, 7, and 14 DPI, p < 0.01 at 4 DPI relative to uninjured fish; remote region: p < 0.05 at 14, 28, and 56 DPI, p < 0.01 at 1 and 4 DPI, p < 0.005 at 7 DPI relative to uninjured fish). Differences between injured and remote regions of the heart at any single timepoint were non-significant, indicating this response is non-localized. Surprisingly, α-smooth muscle actin (αSMA)+/vimentin+ myofibroblasts were not found in any tissue analyzed in this study regardless of injury state or timepoint. The injured zone contained αSMA+ cells at all timepoints beyond 1 DPI in the injured regions of the heart, but αSMA expression did not colocalize with vimentin. This indicates that the myofibroblast phenotype may not be present in the grass carp or may exist in a different state that does not include αSMA upregulation. Together this data indicates widespread vimentin+ fibroblast depletion in the grass carp heart following injury and a lack of traditional myofibroblast differentiation.

**Figure 7:**
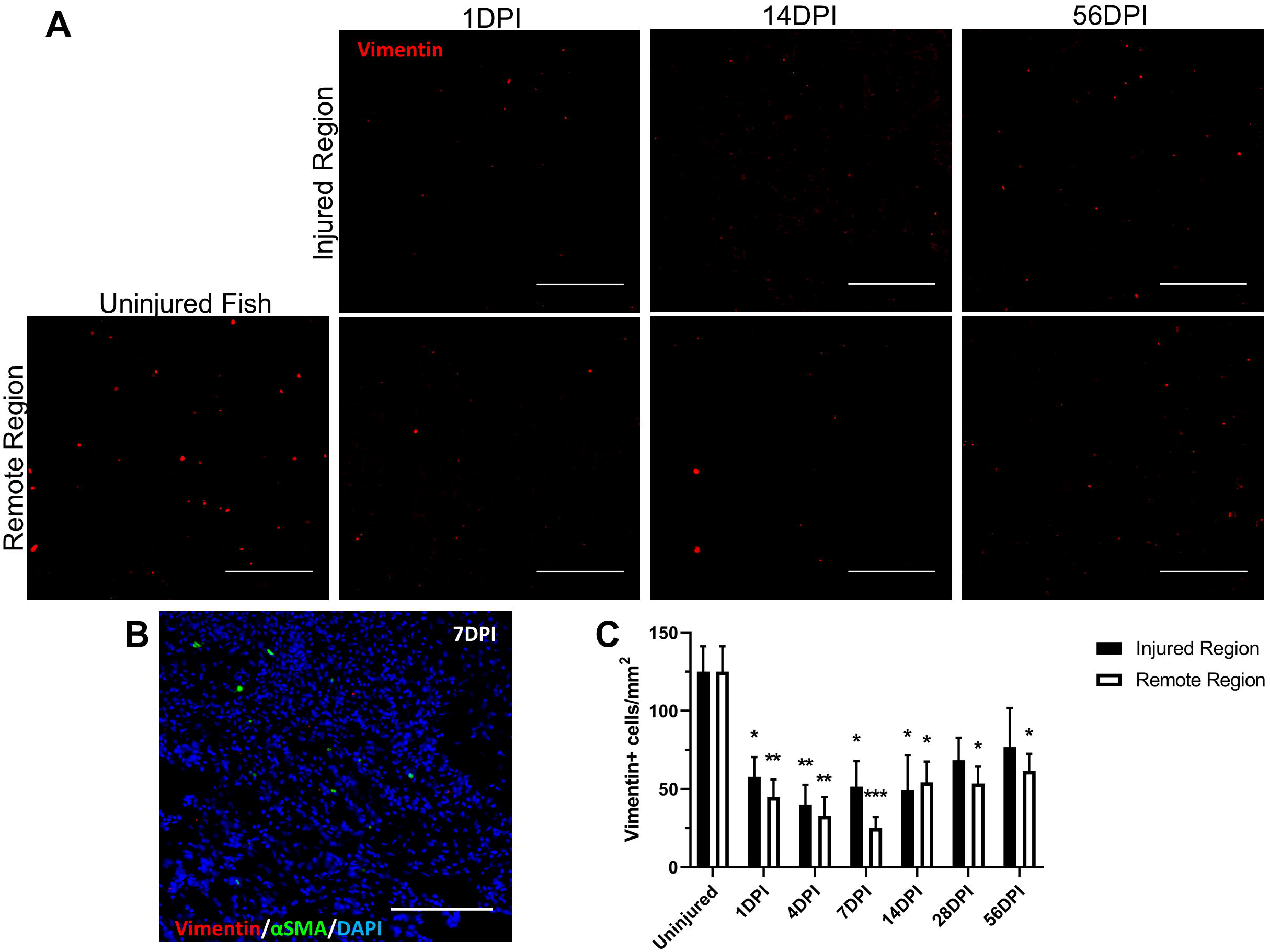
Vimentin+ fibroblasts are depleted following injury. (A) Representative images of vimentin+ fibroblasts are shown in both injured and remote regions of the heart over 56 DPI. (B) aSMA+ cells were present at all timepoints beyond 1 DPI, but no evidence was seen of vimentim+/aSMA+ co-expression, indicating a lack of myofibroblast differentiation in this species following injury (scale bar = 100 μm). (C) Fibroblast density significantly decreases starting and 1 DPI does not fully recover for the remainder of the study. No statistically significant differences are present between injured and remote positions at any time point (scale bar = 100 μm; *p < 0.05, **p < 0.01, ***p < 0.001 relative to uninjured fish).

### 2.7 Macrophage infiltration following cryoinjury

Macrophages are known regulators of cardiac regeneration in zebrafish and fibrosis in non-regenerating species (Aurora et al., 2014; Frantz and Nahrendorf, 2014; Lai et al., 2017). Colony stimulating factor receptor 1+ (Csfr1+) macrophages were stained in both injured and remote regions of the heart following injury to determine their behavior in the grass carp (Chen et al., 2015). Csfr1+ cells were essentially nonexistent in any of the uninjured hearts analyzed. Immediately after injury, csfr1+ cells became more abundant, peaking at 14 DPI before returning to near-baseline levels at 28 and 56 DPI (Fig. 8; injured region: p < 0.05 at 7 DPI, p < 0.001 at 14 DPI relative to uninjured fish; remote region: p < 0.01 at 7 DPI, p < 0.001 at 14 DPI relative to uninjured fish). It is worth noting that csfr1+ macrophages were still present in both injured and remote regions of the heart at 56 DPI (n = 5/5 fish analyzed), although cell densities were not significantly different from baseline. Additionally, macrophage cell density was similar across injured and remote regions of the heart. This indicates that grass carp are neither exhibiting long-term chronic inflammation nor responding specifically to the injured region of the heart following cryoinjury.

**Figure 8:**
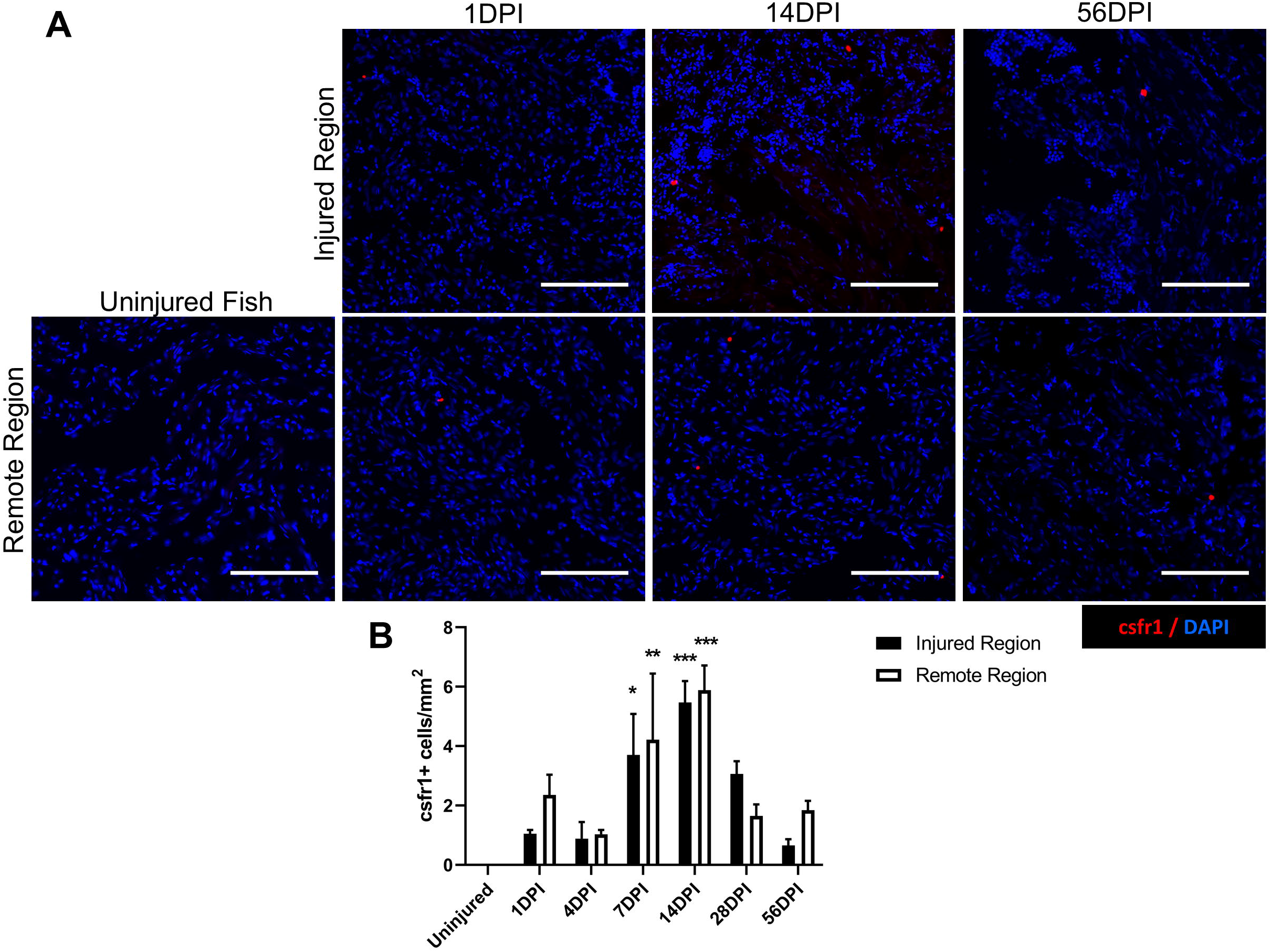
Macrophage populations show a non-localized response to cryoinjury. (A) Representative images of csfr1+ macrophages are shown in both the injured regions and regions of the heart far from the injured zone over time. (B) Quantification of csfr1+ cell density reveals a peak in macrophage density at 14 DPI, although there is no difference in density between injured and remote regions of the heart at any timepoint. (scale bar = 100 μm; *p < 0.05, **p < 0.01, ***p < 0.001 relative to uninjured fish).

## 3. Discussion

Coronary heart disease and its complications remain among the leading causes of death worldwide (Benjamin et al., 2018). Following an injury such as myocardial infarction, the human heart fails to regenerate. Instead, widespread fibrosis and chronic inflammation result in scar formation at the injury site and worsening cardiac function that eventually leads to heart failure (Frantz and Nahrendorf, 2014; Segura et al., 2014; Talman and Ruskoaho, 2016). Naturally, the phenomenon of heart regeneration has gained interest in order to better understand and prevent this adverse remodeling. Several animal models have been studied for this purpose (Cano-Martínez et al., 2010; Flink, 2002; Grivas et al., 2014; Lafontant et al., 2012a; Liao et al., 2017; Neff et al., 1996; Porrello et al., 2011; Poss et al., 2002; Stockdale et al., 2018). Among these models, zebrafish heart regeneration has been most extensively studied and has sparked interest in finding alternate models of cardiac disease in non-mammalian species to determine the driving mechanisms that dictate the cardiac injury response.

Cardiac disease models in teleost fish have focused primarily on the Cypridinae family, which includes zebrafish (Poss et al., 2002), giant danio (Lafontant et al., 2012a), and goldfish (Grivas et al., 2014). Exploration of other fish families has revealed two species of fish that are incapable of cardiac regeneration, the medaka (Ito et al., 2014) and the Pachón variety of *Astyanax mexicanus* (Stockdale et al., 2018). Interestingly, these fish each exhibit unique rather than similar cellular responses to injury that result in chronic fibrosis. While comparisons between these fish and the widely studied zebrafish are possible, the evolutionary distance between these species limits the capabilities of comparative protein or RNA expression techniques (relation between investigated fish are shown in Fig. 1 (Yang et al., 2015)). Additional models of cardiac disease are needed to elucidate the commonalities among species that exhibit chronic fibrosis rather than regeneration. The present study reports the response of grass carp (*Ctenopharyngodon idella*) to cardiac cryoinjury. This is the first reported species within the Cyprinidae family that does not regenerate cardiac tissue after injury. It is also the first large fish species reported in the context of cardiac injury to our knowledge. Instead of widespread cardiomyocyte proliferation and transient fibrosis seen in zebrafish, giant danio, and goldfish (Grivas et al., 2014; Lafontant et al., 2012a; Poss et al., 2002), the grass carp exhibits very limited if any cardiomyocyte proliferation, chronic fibrosis, and decreased cardiac function up to 8 weeks post-injury.

Development of a cardiac injury model in large fish requires additional considerations not relevant to small fish species. The bony operculum in the grass carp restricts access to the ventricle, making resection of the apex impossible, even with the use of retractors. For that reason, a cryoinjury model was developed in which a 2 mm stainless steel probe was applied to the ventricle wall to create a necrotic injured area. This model is not as widely used as resection, but it more closely mimics the initial fibrosis and necrosis seen in human heart disease (Chablais et al., 2011). Unlike small fish species which can be immobilized on a moistened sponge for the short duration of heart injury, the long period of anesthesia required to complete this procedure requires the use of a recirculating pump to allow tricaine anesthetic and water flow across the gills. A recirculation pump system was able to maintain consistent surgical depth of anesthesia with successful recovery from surgery. Additionally, the use of florfenicol antibiotic is a necessary precaution during the injury procedure to prevent the onset of bacterial infection at the incision site, which is not necessary in small fish. Omission of florfenicol resulted in widespread infection and ulceration with 14 days of surgery. We can’t exclude the possibility that florfenicol or Melafix treatments interfere with the carps’ response to cardiac injury as their effect on regeneration has not been documented.

Development of this model ultimately resulted in >95% survival within 14 days of injury and >90% survival to the final 56-day timepoint with consistent injury size that damaged both the compact and trabecular myocardium. High levels of apoptosis were seen as early as 1 DPI in the injured region, which resolved quickly to baseline levels of apoptosis (Fig. 4). This demonstrates the ability of the cryoprobe to create a local and acute injury to the heart that results in an area of tissue necrosis similar to zebrafish and neonatal mouse models (Chablais et al., 2011; Darehzereshki et al., 2015; Polizzotti et al., 2015).

Proliferation of cardiomyocytes is a known driver of cardiac regeneration in non-mammalian species and neonatal mammals (Cano-Martínez et al., 2010; Flink, 2002; Gonzalez-Rosa et al., 2011; Lai et al., 2017; Liao et al., 2017; Porrello et al., 2011; Sallin et al., 2015; Schnabel et al., 2011; Stockdale et al., 2018; Vargas-González et al., 2005; Yu et al., 2018); further, a regenerative phenotype has not yet been reported in the absence of cardiomyocyte proliferation (Ito et al., 2014; Lin and Pu, 2014; Marshall et al., 2017). However, the recent finding of widespread cardiomyocyte proliferation in non-regenerative Pachón *Astyanax mexicanus* suggests that this behavior is not the sole factor that dictates a regenerative response (Stockdale et al., 2018). In the present study, widespread cardiomyocyte proliferation in grass carp was not observed, similar to the response of medaka and adult mammals after injury (Ito et al., 2014; Lin and Pu, 2014).

Proliferation of non-myocytes varies among non-regenerative species, and their underlying functions in the context of cardiac wound healing are currently unclear. Grass carp exhibit an increase in non-myocyte proliferation that peaks at 14 DPI specifically in the injured region (Fig. 4). Interestingly, the non-regenerative Pachón *Astyanax mexicanus* exhibits a similar peak in non-myocyte proliferation at this timepoint, but only in remote regions of the heart rather than the injured zone (Stockdale et al., 2018). Non-regenerative *Xenopus laevis* do not exhibit increased cell proliferation at all (Marshall et al., 2017). Localized proliferation of non-myocytes has thus far only been described in zebrafish and axolotl; however, this occurs at much earlier stages, peaking 1 week post-injury (Chablais et al., 2011; Godwin et al., 2017; Gonzalez-Rosa et al., 2011; Sánchez-Iranzo et al., 2018; Schnabel et al., 2011; Xu et al., 2018; Yu et al., 2018). Taken together, these results indicate that the response of grass carp cardiomyocytes closely resemble non-regenerative medaka and adult mammals, but the proliferation of non-myocytes is a delayed onset of behavior seen in zebrafish during cardiac regeneration. The identification of proliferating non-myocytes in these species has not been fully elucidated but could be key to understanding the species-level differences in cardiac injury response.

Fibrosis occurs in both regenerative and non-regenerative species after cryoinjury due to the presence of necrotic tissue (Chablais et al., 2011; Lai et al., 2017; Marshall et al., 2017; Mizutani et al., 2015; Sánchez-Iranzo et al., 2018). For cardiomyocytes to replace the scar tissue, fibrosis must be transient rather than chronic. Masson’s trichrome staining indicated that scar formation is present in grass carp up to 56 DPI; further, picrosirius red staining revealed that the collagen fibers become more mature and penetrate the trabecular myocardium by 56 DPI, showing no signs of scar regression (Fig. 6). This response is similar to the medaka, where scar formation is seen at 14 DPI (Ito et al., 2014), and *Xenopus laevis*, where scar formation is seen by 1 month post-injury (Marshall et al., 2017). Collagen fibers in zebrafish, however, are visible by 7 DPI, and the scarred region is completely surrounded by healthy tissue by 21 DPI (Chablais et al., 2011). From this comparison, the progression of fibrosis appears similar to the non-regenerative medaka and *Xenopus* laevis.

Surprisingly, grass carp exhibit chronic fibrosis despite a depletion of fibroblasts in the ventricle following injury (Fig. 7). Fibroblasts and myofibroblasts are typically the largest contributors of matrix deposition following injury (Ivey and Tallquist, 2016; Souders et al., 2009) in both regenerative and non-regenerative models. More specifically, fibroblast density in the heart increases within 1 week of injury in every reported species of cardiac injury, including zebrafish (Gonzalez-Rosa et al., 2011; Sánchez-Iranzo et al., 2018; Xu et al., 2018), medaka (Ito et al., 2014), neonatal mice (Mizutani et al., 2015), and adult mammals (Talman and Ruskoaho, 2016). Instead, grass carp see a marked reduction in fibroblasts in the injury zone and beyond that persists across the entire length of the study (Fig. 7). Regardless, scar formation and chronic fibrosis are the outcome in this disease model. These seemingly contradictory results are unique to this species and indicate other cell types are primarily responsible for secreting fibrotic ECM within the injured region. In support of this theory, zebrafish endocardial cells are known to secrete collagen within the injury region 7 DPI (Sánchez-Iranzo et al., 2018), although their ultimate contribution to scar formation is limited relative to fibroblasts. Endocardial cells may be playing a role in grass carp scar formation as well, but we were unfortunately unable to assess the cell type contributing to this fibrotic behavior due to availability of species-specific reagents. Future studies could identify the role of endocardial cells in scar formation and maintenance in the injured grass carp heart. Another surprising result in this species is the lack of αSMA+/vimentin+ myofibroblasts (Fig. 7B). Other studies have used αSMA as a putative marker of myofibroblasts and shown infiltration of these cells to the injured region (Ito et al., 2014; Yu et al., 2018), but the lack of colocalization shown in this study indicate that αSMA alone is not sufficient to identify myofibroblasts in fish. Our data indicates that myofibroblasts, if they exist in the grass carp, could have a different expression profile than mammals, making the colocalization of αSMA and vimentin an ineffective probe for myofibroblast differentiation.

The immune response has repeatedly been shown as a vital component of regeneration. In zebrafish, depletion of macrophages impedes regeneration (Lai et al., 2017), while stimulation of macrophages in the non-regenerative medaka promotes cardiomyocyte proliferation and scar resolution (Lai et al., 2017). Grass carp exhibit a delayed and muted csfr1+ macrophage response, peaking at 14 DPI; in contrast, macrophage infiltration peaks between 1 and 4 DPI in zebrafish and *Xenopus tropicalis* and reach baseline by 14-16 DPI (De Preux Charles et al., 2016; Liao et al., 2017; Xu et al., 2018). Macrophage presence in the grass carp does not mimic the chronic response seen in mammals either. Adult mice exhibit a macrophage peak near 5 DPI (Talman and Ruskoaho, 2016) and remain above baseline for at least 8 weeks (Sager et al., 2016) while grass carp reach near-baseline levels by 28 DPI. Surprisingly, macrophages are not restricted to the injured region, and an organ-wide response is seen instead (Fig. 8). A previous study in the zebrafish and medaka suggests that early macrophage response is essential to a regenerative phenotype, so this finding at least partially explains the lack of regeneration seen in the grass carp (Lai et al., 2017). However, the effect of a localized versus organ-wide immune response has not yet been explored and warrants further investigation.

Due to the novelty of this model in the context of cardiac disease, caveats of using the grass carp as a study model must be acknowledged. The size of these fish relative to other fish models imposes restrictions on the housing and throughput capabilities in a laboratory setting. The injury protocol takes longer in grass carp than in zebrafish and has the potential to cause higher levels of stress. A previous study in zebrafish has examined the detrimental effects of stress on cardiac regeneration (Sallin and Jaźwińska, 2016), and stress may have exasperated the fibrotic response seen in the present study. Due to the sparsity of grass carp studies performed, molecular techniques and protein characterization in this species is not as well-developed as other models of cardiac disease such as mice and zebrafish. This limits the ability to identify specific cell types or proteins in the heart. However, with recent publication of the grass carp genome (Wang et al., 2015) and widespread availability of total protein and gene expression techniques, this limitation can be addressed in future studies.

The present study, in addition to identifying a new potential model of cardiac disease, describes a surgical procedure for cardiac injury adaptable to other large species of fish. Although there are essentially two phenotypes to cardiac injury, our evidence in the grass carp along with the contributions of others suggest that a fibrotic rather than a regenerative response can be due to multiple different changes in cell behavior. While Pachón *Astyanax mexicanus*, grass carp, and medaka all fail to regenerate, multiple differences in cell behavior exist between these species that each lead to similar fibrotic phenotypes. The grass carp provides a model to study fibrosis apparently independent of fibroblast infiltration. Future studies in these animals could identify cell types and proteins previously unknown to contribute to chronic fibrosis in the context of cardiac disease. Studying these models has implications not only to understanding the underlying mechanisms of regeneration and chronic fibrosis but also in identifying potential targets to treat human heart disease.

## 4. Materials and Methods

### 4.1 Animal Procurement and Care

All work with live animals was approved by the Cornell University Institutional Animal Care and Use Committee prior to starting experiments. Results from a total of 35 fish are presented in this study. Live grass carp weighing 435 ± 112 g were purchased from a local fish farm (Candor, NY, USA) (Fig. 1B). Fish were housed in indoor 360 gallon capacity tanks measuring roughly 1.2 × 1.2 × 0.9 meters with at least once full water turnover a day. Water temperature was kept between 18 and 20°C. Nets and tarps over the tanks prevented fish from jumping out of the tank, a natural defense mechanism in this species. A maximum of 6 fish were housed per tank. Fish were fed commercial trout pellets several times per day. Any fish showing signs of fin rot were isolated and treated with 7-day soaks in aquarium salt (1.5 tsp/gal) and a 7-day treatment with Melafix (API Fishcare, Chalfont, PA, USA) according to package instructions.

### 4.2 Cryoinjury procedure

To induce anesthesia, carp were captured individually and submerged in a bath of tricaine methanesulfonate (MS-222) and sodium bicarbonate (both at 150 mg/L, tricaine supplied by Western Chemical Inc., Ferndale, WA, USA) to induce anesthesia. Surgical depth anesthesia was observed by slowed or stopped opercula movements, ceased body movements, lack of response to a pinch at the base of the tail, and floating in a supine position. A recirculating water pump was then fitted to the fish’s mouth to continuously flush a tricaine and sodium bicarbonate solution (each at 100 mg/L) over the gills for the duration of surgery. After placing the fish in a supine position, excess mucus was removed from the underside with gauze and the surgical surface was cleaned twice with povidone-iodine applied in the direction of scale growth. Pre-surgical analgesia of lidocaine (Henry Schein Animal Health, Dublin, OH, USA) was administered subcutaneously (1 mg/kg), and carprofen (Putney Inc., Portland, ME, USA) was administered IM (2.5 mg/kg); a dose of florfenicol (30mg/kg; Merck Animal Health, Summit, NJ, USA) was administered IM to prevent bacterial infection due to surgery.

Three to four rows of scales directly overlying the heart were removed by pulling caudally until they released. The exposed skin was washed twice with povidone-iodine solution. The heart was exposed through a 1.5 cm incision along the length of the fish, cutting through the skin, underlying musculature, and pericardial sac. Alm retractors (17008-07; Fine Science Tools Inc., Foster City, CA, USA) were used to separate the operculum approximately 5-8 mm, allowing direct visualization of the ventricle (Fig. 1C). Excess blood and moisture were removed from the ventricle surface and body cavity with cotton-tipped swabs. A liquid nitrogen-cooled stainless steel probe measuring 2 mm in diameter was then directly applied to the ventricle for 30 seconds along the free wall as close to the apex as possible to create the cryoinjury. After 30 seconds, a few drops of saline were used to thaw the probe and tissue, allowing trauma-free removal of the probe. A successful injury is easily observed due to an instantaneous lack of ventricle movement that occurs upon application of the cryoprobe and recovers quickly after thawing. Additionally, the injured region is seen as a bright red mark on the ventricle surface after removing the cryoprobe. Excess moisture and blood were removed from the body cavity and ventricle surface before removing the retractors. The incision was closed using 4-0 polyglyconate suture in a simple interrupted pattern, closing the muscle and skin layers separately whenever possible. Immediately after wound closure, the fish was moved to a recovery tank and fresh water moved constantly over its gills until opercular movements and swimming behavior recovered. Following recovery, the fish was returned to its original tank.

### 4.3 Measuring cardiac function via echocardiography

Immediately before sacrifice, fish were anesthetized in a tricaine/sodium bicarbonate bath (each at 150mg/L) and then maintained at 100mg/L. A Vevo 2100 series ultrasound fitted with a MS400 transducer (FUJIFILM VisualSonics Inc., Toronto, ON, Canada) was used to obtain long-axis images of the heart in at least 3 separate locations. Short-axis imaging was not possible due to interference by the operculum. Immediately after imaging, the fish was placed in a high concentration tricaine/sodium bicarbonate bath (each at 300mg/L) for euthanasia.

FAC, a functional measurement of cardiac function, was calculated by manually outlining and measuring the area occupied by the ventricle at systole and diastole. FAC percentage was then calculated as 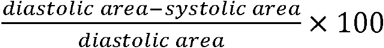. FS, another measurement of ventricular contractility, was determined by measuring the distance from base to apex at systole and diastole. Similarly, FS, expressed as a percentage, was calculated as 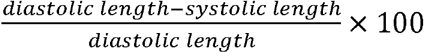. At least 3 separate heart cycles were analyzed for FAC and FS measurements. Echocardiography-based measurements were made using ImageJ (NIH, Bethesda, MD, USA). Decreases in either of these measurements are indicative of a loss in cardiac function, and both are used clinically to assess heart function after myocardial infarction (MI).

### 4.4 Tissue processing and histology

Immediately after euthanasia, the entire heart was extracted, including the bulbous arteriosus and atria. The atrium was often destroyed during processing and is not always present in histology samples. Hearts were fixed for 1 hour in 4% (w/v) paraformaldehyde, cryoprotected overnight in 30% (w/v) sucrose, embedded in O.C.T. compound, and frozen in dry ice-cooled isopentane. Long-axis serial sections were collected at a thickness of 8 µm with each section separated by roughly 400 µm on the slide.

AFOG staining was performed as previously described (Darehzereshki et al., 2015; Poss et al., 2002) to mark healthy tissue, fibrin, and collagen. For each of these stains, injury area was calculated as percentage of the ventricle area covered by collagen or fibrin-stained tissue. This was repeated for each tissue section, and the largest relative injured area was taken as the injury size for that heart.

Masson’s trichrome staining was performed to indicate which timepoints following injury were characterized by collagen deposition and therefore appropriate for picrosirius red staining. Picrosirius red staining was then performed on these samples to assess the maturity and alignment of collagen fibers within the ventricle after injury under polarized light microscopy. Within the injured region of each heart, the distribution of green, yellow, orange, and red fibers were calculated using MATLAB as previously described (Rich and Whittaker, 2005). Immature collagen fibers were defined as green and yellow fibers, and mature fibers were defined as orange and red. The percentage of mature fibers in the injured region was used for statistical comparisons, although data for all defined colors are presented. Alignment was calculated as previously described (Long et al., 2017) using a fast Fourier transform (FFT) macro in ImageJ, where index values close to 1 indicate random fiber alignment and higher index values indicate increased fiber alignment.

### 4.5 Apoptosis in the heart following injury

To assess levels of apoptosis in carp hearts following cryoinjury, the Click-iT™ Plus TUNEL assay (Invitrogen, Carlsbad, CA, USA) was used according to the manufacturer’s instructions. Prior to staining, tissue sections were fixed with 4% formaldehyde for 15 minutes followed by permeabilization in 0.25% (v/v) triton x-100 for 15 minutes. The percentage of apoptotic cells was assessed in both the injured region and an uninjured region of the heart far from the injury (remote) to determine the role of both local and systemic cell death in the tissue response.

### 4.6 Immunofluorescent staining and imaging

Immunofluorescent staining of tissue sections was used to determine the behavior of different cell populations following cryoinjury. All immunofluorescent stains were performed after 5-minute fixation in 2% (w/v) formaldehyde, permeabilizing for ten minutes in 0.2% (v/v) triton x-100, and blocking for 1 hour in 5% normal goat serum at room temperature. Primary antibodies were incubated in 1% normal goat serum at 4ºC overnight and detected with fluorescent secondary antibodies (A-11032, A-11037, A-11029, ThermoFisher Scientific, 1:500 dilution) through a 1-hour incubation at 37ºC. All brightfield and immunofluorescent imaging was performed on a Nikon Eclipse Ti2-E inverted microscope, and image analysis performed with NIS Elements software unless otherwise specified (Tokyo, Japan).

Proliferation of cardiomyocytes is a known mechanism of cardiac regeneration in zebrafish (Chablais et al., 2011; Jopling et al., 2010; Schnabel et al., 2011). Therefore, the presence of actively proliferating cardiomyocytes was assessed by co-staining of cardiac troponin T (cTnT; MA5-12960, ThermoFisher Scientific, 1:40 dilution) found in cardiomyocytes and pH3 (ab5176, Abcam, 1:200 dilution), a mitotic marker. The density of pH3+ cardiomyocytes and pH3+ non-myocytes were calculated at each timepoint within the injured region and a remote region of the ventricle.

Fibroblasts and myofibroblasts are responsible for secreting and remodeling the extracellular matrix (ECM) following cardiac injury (Ma et al., 2017). The density of vimentin+ cells (ab8069, Abcam, 1:100 dilution) in injured and remote regions were quantified over time, and myofibroblasts, defined as vim+/αSMA+, (αSMA, ab8211, Abcam, 1:100 dilution) were identified to detect a potential pro-fibrotic response. Immune cell response following injury is known to play an active role in cardiac regeneration in other species (Lai et al., 2017; Xu et al., 2018). Therefore, csfr1+ (GTX128677, Genetex, 1:200 dilution) macrophage density was assessed by immunofluorescent staining of both injured and remote regions of the grass carp ventricle following injury to detect both the location and time-dependence of macrophage infiltration into the damaged tissue.

### 4.7 Data and statistical analysis

All data is presented as mean ± SEM, and n = 3 – 5 fish per timepoint were used for each measurement. Statistically significant differences in means were detected using independent analysis of variance (ANOVA); one-way independent ANOVA was used when a single measurement was made per animal, and two-way independent ANOVA was used when comparisons were made across time and between remote and injured regions of the heart. Tukey’s post hoc testing was used for AFOG injury size, proliferation of non-myocytes, and picrosirius red fiber color distribution and alignment in order to compare across all timepoints and regions. Sidak’s post hoc test was used to compare FAC, FS, TUNEL, fibroblast density, and csfr1+ macrophage density measurements relative to uninjured fish samples.

## 5. Acknowledgements

We thank Dr. Kenneth Poss and Dr. Voot Yin for providing their AFOG protocols and the entire veterinary and animal care staff at Cornell University for their assistance throughout this study. Additionally, we thank Dr. Helene Marquis for assisting with our fish anesthesia systems and monitoring.

